# A metagenomic library cloning strategy that promotes high-level expression of captured genes to enable efficient functional screening

**DOI:** 10.1101/2023.03.24.534183

**Authors:** Michelle H Rich, Abigail V Sharrock, Timothy S Mulligan, Frazer Matthews, Alistair S Brown, Hannah R Lee-Harwood, Elsie M Williams, Janine N Copp, Rory F Little, Jenni JB Francis, Claire N Horvat, Luke J Stevenson, Jeremy G Owen, Meera T Saxena, Jeff S Mumm, David F Ackerley

## Abstract

Functional screening of environmental DNA (eDNA) libraries is a potentially powerful approach to discover enzymatic “unknown unknowns”, but is usually heavily biased toward the tiny subset of genes preferentially transcribed and translated by the screening strain. We have overcome this by preparing an eDNA library via partial digest with restriction enzyme FatI (cuts CATG), causing a substantial proportion of ATG start codons to be precisely aligned with strong plasmid-encoded promoter and ribosome-binding sequences. Whereas we were unable to select nitroreductases from standard metagenome libraries, our FatI strategy yielded 21 nitroreductases spanning eight different enzyme families, each conferring resistance to the nitro-antibiotic niclosamide and sensitivity to the nitro-prodrug metronidazole. We showed expression could be improved by co-expressing rare tRNAs and encoded proteins purified directly using an embedded Hisg-tag. In a transgenic zebrafish model of metronidazole-mediated targeted cell ablation, our lead MhqN-family nitroreductase proved ∼5- fold more effective than the canonical nitroreductase NfsB.

## Introduction

Bacteria-derived enzymes have innumerable applications in research, medicine and industry, with the industrial enzymes market alone projected to exceed US$10 billion by 2024 (Berini et al, 2017). However, an overwhelming majority of bacteria cannot be cultivated efficiently in the laboratory, meaning that traditional microbiological methods are limited in their scope to screen for new enzymes with desirable activities (Schmeisser et al, 2007; Uchiyama and Miyazaki, 2009; Berini et al, 2017). To address this, culture-independent strategies have been developed to discover enzymes encoded by metagenomic DNA, extracted from promising environments. Frequently these approaches are sequence-based, employing next-generation sequencing and bioinformatics to identify candidate genes (Schmeisser et al, 2007; Berini et al, 2017). Although these approaches can traverse a vast amount of sequence space, the need to subsequently synthesise or amplify, clone and characterise each candidate is a major bottleneck that adds substantial time and expense. Moreover, sequence-based approaches are limited by their inherent need for similarity. Not only can highly divergent homologues of known enzymes be difficult to identify (Bernard et al, 2018), microbial metagenome sequencing is revealing ever-increasing numbers of hypothetical proteins of unknown function that can nevertheless be highly effective at catalysing a desired reaction (Vanni et al, 2022).

An alternative is functional screening, whereby metagenomic DNA fragments are cloned in a vector, expressed in a host strain such as *Escherichia coli,* and the desired activity is screened or selected at a phenotypic level (Schmeisser et al, 2007; Uchiyama and Miyazaki, 2009; Ngara et al, 2018). This avoids sequence-level preconceptions, but can introduce other significant biases, most notably that the host cell is often unable to effectively recognise transcriptional and/or translational signals from evolutionarily distant species, resulting in low-to-no expression of most captured genes (Uchiyama and Miyazaki, 2009; Han et al, 2022). Functional metagenome screening also requires an effective high-throughput screen, or ideally a selection, to recover clones that gain the desired activity (Bunzel et al, 2018; Ngara et al, 2018; Markel et al, 2020).

We have a strong interest in discovery and characterisation of bacterial nitroreductases that can efficiently convert prodrugs to a toxic form, by reducing an electron-withdrawing nitro substituent on an aromatic ring to an electron-donating hydroxylamine or amine via concerted two-electron transfer steps (Williams et al, 2015). Nitro-reduction is a function-based classification that encompasses diverse enzyme families and, given the paucity of nitroaromatic molecules in nature (Parry et al, 2011), is generally assumed to be a promiscuous activity (Roldán et al, 2008; Hall et al, 2020). Known nitroreductases include members of the NfsA, NfsB, PnbA quinone oxidoreductase families that share a conserved “nitroreductase” structural fold (Akiva et al, 2017), but also of the AzoR (azoreductase), MsuE (sulphur assimilation) and NemA (old yellow enzyme) families, which share little sequence or structural homology other than all binding FMN or FAD cofactors (Green et al, 2013; Prosser et al, 2013). There are undoubtedly many other families of potential nitroreductases that remain to be discovered, and even within the known enzyme families it is difficult to predict *a priori* whether a given member is likely to be an efficient nitroreductase. Thus, functional metagenome screening is an attractive strategy to discover new nitroreductases.

Our primary focus here was discovery of nitroreductases that can efficiently convert the nitro­prodrug antibiotic metronidazole to a cytotoxic form. Metronidazole has been used to precisely ablate target cells in transgenic model organisms that express the *E. coli* nitroreductase gene *nfsB* from a cell-type-specific promoter, for the purposes of investigating cellular function and/or regeneration (Curado et al, 2007). This system has received widespread uptake, in particular in zebrafish, but is confounded by the need for metronidazole concentrations near the toxicity threshold (~10 mM) to achieve effective ablation of many cell types (White et al, 2013; Mathias et al, 2014). More efficient metronidazole-converting nitroreductases would therefore enable improved ablation with fewer off-target effects. To select for metagenome-derived nitroreductases we hoped to use niclosamide, an antibacterial that we previously found is not only detoxified by nitro­reduction, but was also able to select for more effective metronidazole-reducing variants of the *E. coli* nitroreductase NfsA from targeted mutagenesis libraries (Copp et al, 2020; Sharrock et al, 2021). An important caveat is that those previous mutagenesis studies employed high-level expression of each nitroreductase gene variant from a strong *tac* promoter on a high copy number plasmid. We were therefore uncertain whether niclosamide would prove effective in recovering nitroreductases from a metagenome library, which were likely to be expressed at far lower levels. Ultimately, we found it necessary to implement an alternative strategy that minimises species bias effects and promotes high-level expression of captured genes. This strategy is likely to be broadly applicable for discovery of any catalytic functionality for which an effective screen or selection can be applied.

## Results

### Niclosamide can select for metronidazole-active nitroreductases

Although niclosamide is usually far more toxic to gram-positive than gram-negative bacteria (Rajamuthiah et al, 2015; Peyclit et al, 2022), we have previously shown that deletion of the *tolC* efflux gene and seven endogenous nitroreductase genes renders *E. coli* 2000-fold more sensitive to niclosamide (Copp et al, 2020). The majority of this sensitisation effect was due to loss of *tolC,* but we have nevertheless shown that over-expressed mutants of the *E. coli* nitroreductase *nfsA* provide a selectable level of niclosamide resistance in this multi-gene deletion strain (F. *coli* 7NT; Copp et al, 2020; Sharrock et al, 2021). To test whether the ability to detoxify niclosamide is widely associated with sensitivity to metronidazole, we measured the growth of 18 7NT strains individually over­expressing members of an oxidoreductase gene library (representing six known nitroreductase families) in lysogeny broth amended with 0.8 µM niclosamide or 800 µM metronidazole. Although there was only a moderate inverse correlation (r^2^ = 0.43; Pearson’s r = -0.65) between levels of growth in each medium, the data were skewed by three AzoR family members that conferred high levels of niclosamide resistance, but little sensitivity to metronidazole **(Supplementary Figure S1).** Overall, 10 of the 13 strongly niclosamide-resistant strains were overtly growth-inhibited by metronidazole, compared to none of the niclosamide-sensitive strains.

### Niclosamide is inefficient in selecting nitroreductases from standard metagenome libraries

*We* next conducted pilot tests to assess whether niclosamide could efficiently select for nitroreductase genes from a typical metagenome library, i.e. one generated by purification, fragmentation and cloning of environmental DNA (eDNA) in a standard *E. coli* expression vector. For this, we used a small and well-characterised soil eDNA library containing *ca.* 1.3 × 10^5^ unique metagenome inserts with an average size ~4 kb, cloned into pRSETB (Parachin and Gorwa-Grauslund, 2011). In an earlier study we used this library to functionally screen for 4’-phosphopantetheinyl transferase genes as markers for natural product biosynthetic gene clusters, and recovered seven unique inserts (Owen et al, 2012). Bacterial genomes typically encode numerous nitroreductases apiece (Prosser et al, 2013; Akiva et al, 2017), so we considered that recovery of >10 unique nitroreductases would indicate an efficient selection.

As all nitroreductase genes from our oxidoreductase library **(Supplementary Figure S1)** and in our previous niclosamide resistance screens (Copp et al, 2020) had been strongly over-expressed, we sought to boost transcription of insert DNA from the T7 promoter of pRSETB via IPTG induction (as had been used for this eDNA library by Parachin and Gorwa-Grauslund, 2011). For this, we first lysogenised the 7NT strain with *λ*DE3, which carries a T7 RNA polymerase gene. The resulting *E. coli* strain (7TL) was transformed with the soil eDNA library, and selection with 0.5 µM niclosamide (the lowest concentration that reliably prevented colony formation by empty plasmid control cells) yielded 21 niclosamide-resistant colonies. While this initially appeared promising, Sanger sequencing revealed that these 21 ‘hits’ represented only three unique inserts, each of which contained a *tolC-*like gene **(Supplementary Table S1).** Resistance was therefore likely due to restoration of efflux rather than detoxification of niclosamide.

TolC-mediated efflux of niclosamide can be prevented by the chemical inhibitor phenylalanine-arginine-β-naphthylamide (PAβN; Copp et al, 2020), so we added PAβN to the selection medium and re-screened the library. However, we did not recover any eDNA clones at concentrations of niclosamide and PAβN that prevented growth of an empty plasmid control strain while permitting colony formation by 7TL cells that expressed *E. coli* NfsB from a *tac* promoter. We therefore concluded that niclosamide did not provide an efficient means to select nitroreductase genes from standard eDNA libraries. It seemed likely that this was because nitroreductase-mediated niclosamide resistance requires higher-level expression of captured genes than a standard eDNA library can routinely provide (whereas trace levels of *tolC* expression appeared sufficient to confer resistance).

### A Fatl eDNA cloning strategy for selection of niclosamide and metronidazole active nitroreductases

An ideal solution to boost gene expression would be to ligate eDNA fragments into a plasmid in such a way that the start codon of a captured gene was placed an optimal distance downstream of a strong promoter and ribosome binding site. In considering this problem, we realised that the most common bacterial start codon (ATG) constitutes three quarters of the palindrome recognised by the restriction enzyme *Fatl* (^↓^CATG). We envisaged that a partial *FatI* digest of eDNA would yield an array of fragments with 5’ overhangs that often contain start codons, allowing their associated genes to be ligated into a custom expression vector at a precise location. We therefore designed a plasmid with a unique and compatible *NcoI* site (C^↓^CATGG) located downstream of an IPTG-inducible *tac* promoter and strong ribosome binding site (an inducible promoter was chosen to avoid prematurely selecting against gene inserts that impose a fitness burden upon the host cell). Reasoning that it might sometimes be useful to purify target proteins directly from selected bacterial clones, we also embedded an optimally-positioned start codon and N-terminal hexahistidine tag directly upstream of, and in frame with, the ATG of the captured gene (**Figure** 1). The final plasmid (pUCXMG; **Supplementary Figure S2**) was assembled from an artificially-synthesised DNA fragment ligated into a pUCX parental plasmid backbone. Pilot tests demonstrated that a nitroreductase gene *(azoR)* cloned into the *NcoI* site of pUCXMG conferred a similar level of *E. coli* host cell protection against niclosamide to *azoR* expressed from the parental pUCX plasmid **(Supplementary Figure S3**).

**Figure 1:**
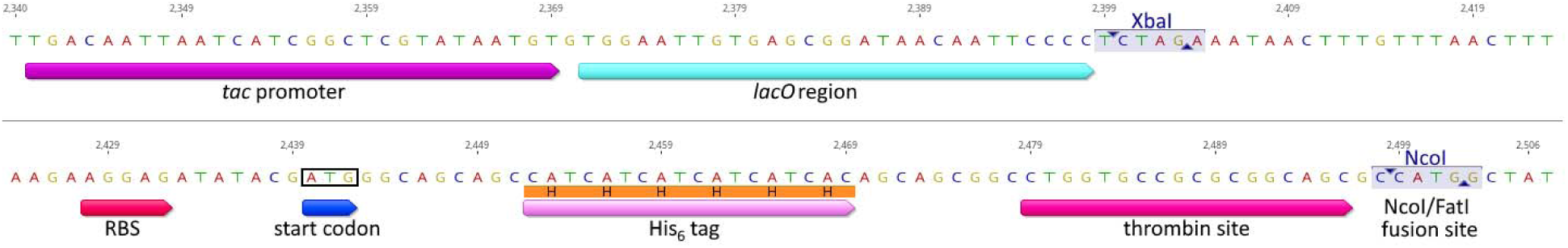
Key features of FatI expression vector pUCXMG. Highlighted are the IPTG-inducible *tac* promoter; the *lacO* operator (repressor-binding) region; an XbaI site used in vector assembly; a strong ribosome binding sequence (RBS; derived from the T7 phage major capsid protein RBS); the start codon; the hexahistidine (His_6_) tag; a thrombin cleavage sequence for His_6_ tag removal; and the NcoI restriction site for insertion of FatI partially-digested eDNA fragments. Figure drawn using Geneious Prime version 2022.2. The full plasmid map and sequence are available as Supplementary Figure S2.

The *Fatl* partial digest strategy we envisaged only permits precision cloning of the subset of genes that possess both an ATG start codon and a cytosine in the -1 position; but soil eDNA is such a vast resource that we did not consider this a significant limitation. Nevertheless, we felt it important to consider the distribution of genes that possess these characteristics, for example, to assess the extent to which our method might bias for genes from GC-rich bacteria. For this, we collected 21,675 annotated bacterial genomes from the National Centre for Biotechnology Information (NCBI) Assembly Database, and wrote a Python script to analyse the number of genes using each start codon (ATG, GTG, TTG) and the corresponding nucleotide distribution at the -1 position, within each genome. To exclude plasmid sequences from the analyses, we performed analyses on records within each genome without ‘plasmid’ in the record description; and for genomes containing multiple chromosomes, the results from all chromosomes were combined into a single record (see **Supplementary Files S1 and S2** for bioinformatics scripts and compiled genome analyses). When we plotted the proportion of ATG and (C)ATG start codons (where () denotes the nucleotide in the -1 position) in each genome relative to its GC content, we noticed a disproportionately high incidence of (C)ATG start codons relative to (G)ATG (**Figure 2**). Indeed, in a substantial proportion of bacteria containing >60% GC content, over 50% of genes initiate with a (C)ATG start codon. This is helpful from the perspective of capturing coding sequences effectively, but does suggest that DNA from these species will be overrepresented in metagenomic libraries prepared via *FatI* partial digestion.

**Figure 2:**
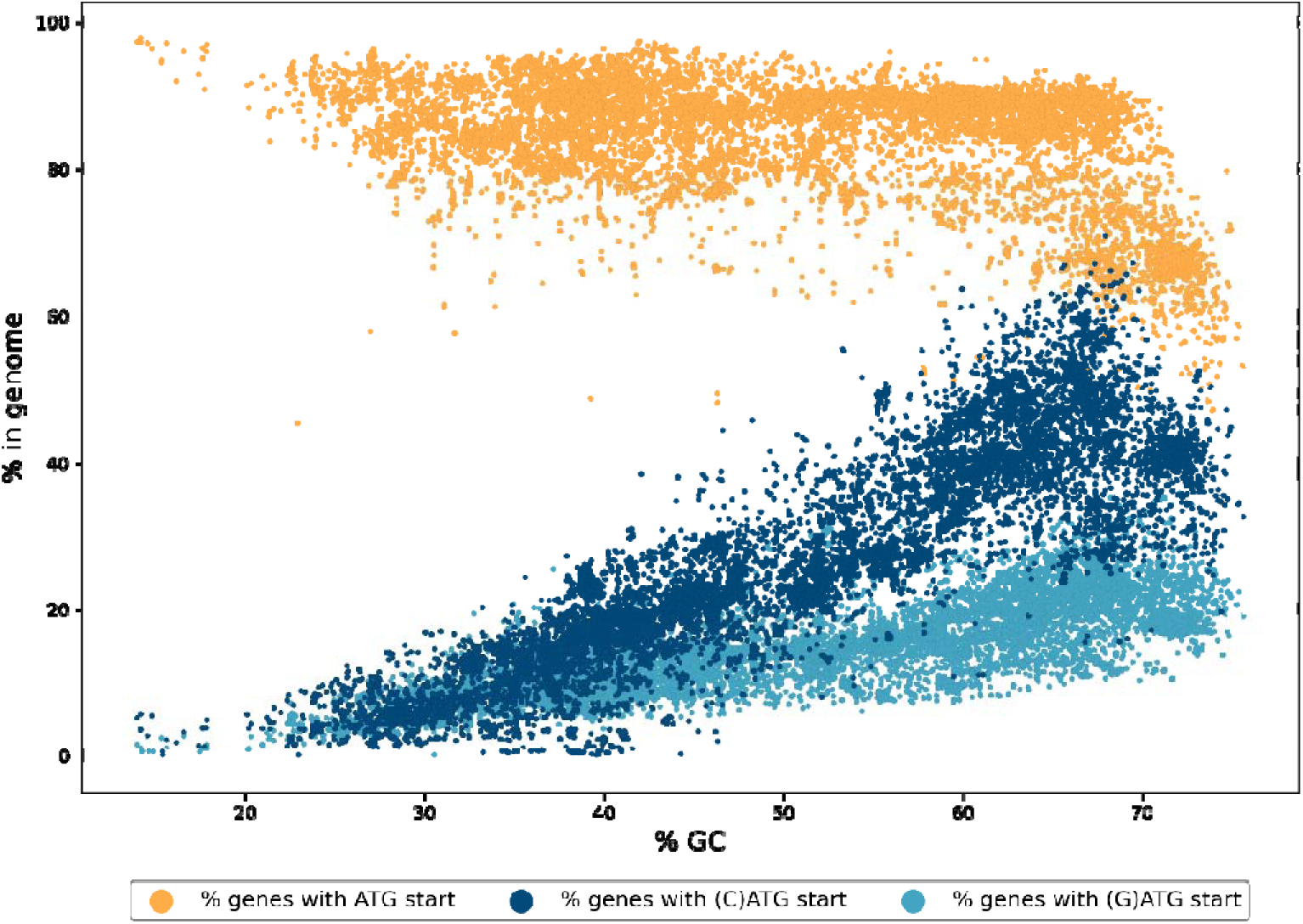
Percentage of genes from sequenced genomes that initiate with ATG, (C)ATG or (G)ATG start codons, relative to genomic GC content. The percentage of genes predicted to initiate with ATG (orange), (C)ATG (dark blue), or (G)ATG (light blue) start codons were sourced from 21,675 annotated bacterial genomes, derived from the National Centre for Biotechnology Information Assembly Database on June 7, 2021, and plotted relative to the total GC content of that genome. Data were analysed with Python 3.8.1, using Script 1 (Supplementary Files).

To implement our cloning strategy (**Figure** 1), we purified DNA from 250 g of locally-collected soil. This yielded 38 µg of purified DNA that was primarily of a size range >10 kb **(Supplementary Figure S4**). Following partial digestion with *FatI* **(Supplementary Figure S4**), we gel-extracted DNA fragments in the 0.6-1.4 kb range, seeking to (i) emphasise single-gene inserts that are more amenable to ligation, one-pass Sanger sequencing and deconvolution of phenotypes; and (ii) capture a wide diversity of bacterial nitroreductases while excluding *tolC* genes (typically >1.5 kb). Upon ligation of these fragments into the *NcoI* site of pUCXMG, we generated a plasmid library of 1.38 × 10^7^ clones, with an estimated insert rate of 87.5% i.e. 1.2 × 10^7^ unique variants in total (~1.0 × 10^7^ with an insert >500 bp; **Supplementary Figure S5**).

In two independent experiments, *E. coli* 7TL cells transformed with this library were plated to an estimated 10-fold coverage on niclosamide-amended media. In total, 910 resistant colonies were selected and then counter-screened for host-cell sensitivity to 1.5 mM metronidazole (**Figure 3**). This yielded 178 metronidazole-sensitive ‘hits’ that were sent for Sanger sequencing of the plasmid insert, revealing 21 unique inserts. Each of these contained a gene predicted (by BLAST alignment) to encode a flavin-associated protein, a substantial majority (17/21) of which were ligated in-frame at the *Ncol/Fatl* fusion site of their recombinant pUCXMG vector (**Table** 1). To eliminate possible chromosomal mutations, each unique plasmid was used to transform fresh *E.* co//cells, and the resulting strains were then subjected to quantitative IC_50_ assays (**Table** 1). Nine of the 21 recovered nitroreductases were found to sensitise *E. coli* host cells to lower concentrations of metronidazole than *E. coli* NfsB, the benchmark enzyme for metronidazole-mediated cell ablation. The most active enzyme (MhqN1) sensitised *E. coli* to a 24-fold lower metronidazole concentration and was identified through BLAST analysis as belonging to the MhqN clade of the ‘nitroreductase superfamily’ (Akiva et al, 2017). The second most-active enzyme (MhqN2) also belonged to the MhqN family (Table 1).

**Figure 3:**
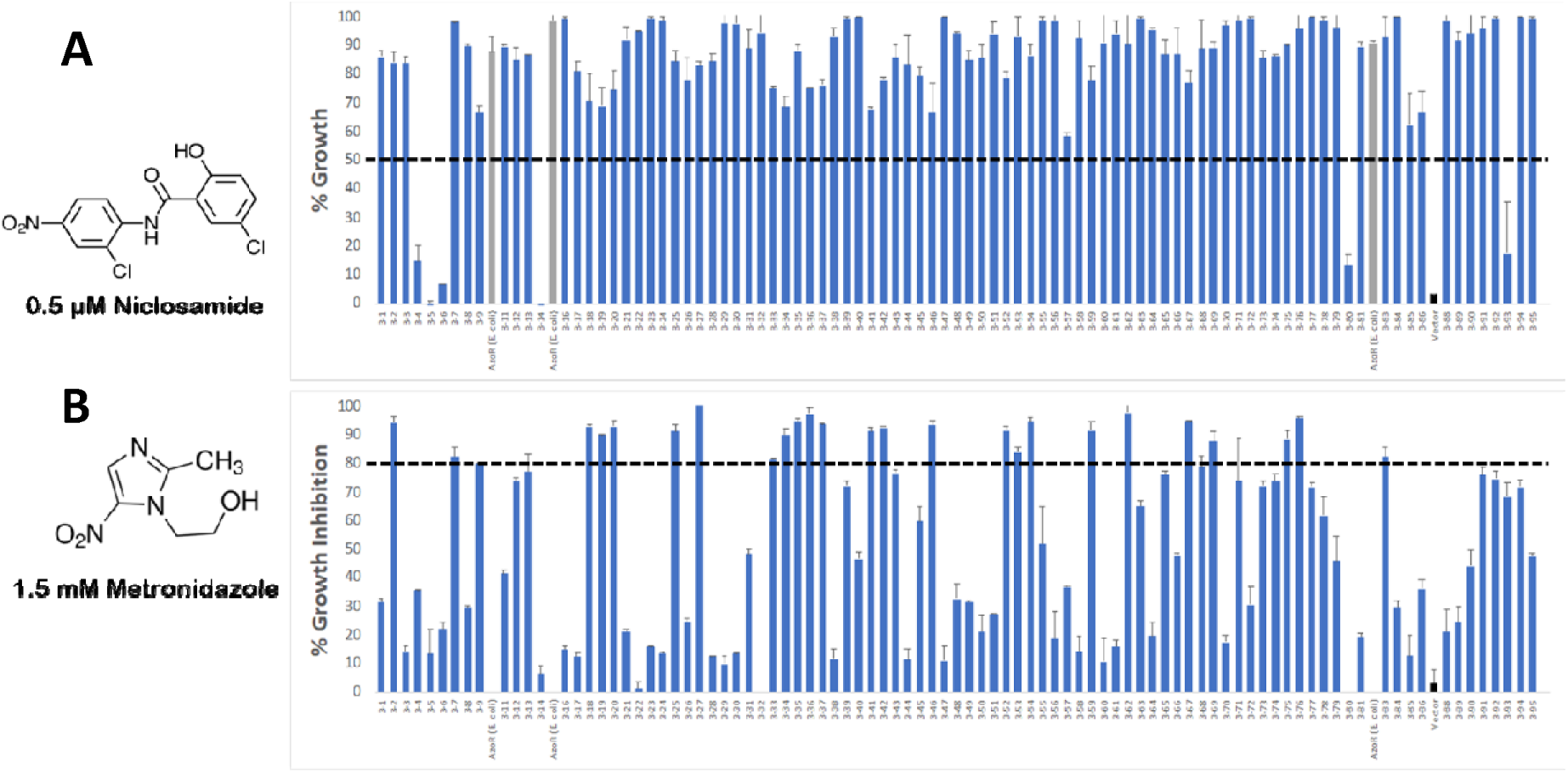
Counter-screening of niclosamide-resistant *E. coli* 7TL eDNA variants to identify metronidazole sensitive strains. 910 niclosamide-resistant colonies were recovered from plating of *E. coli* 7TL cells transformed with the FatI eDNA library on LB agar amended with 0.5 µM niclosamide. Replicate LB cultures were established from each colony and grown for 4 h in either unamended media as a control or else media amended with 0.5 µM niclosamide or 1.5 mM metronidazole. The percentage growth of niclosamide-challenged cultures **(A)** or percentage growth inhibition of metronidazole-challenged cultures **(B)** were calculated relative to the unchallenged control. Panels A and B present data from a single set of representative 96-well plates (each of which contained a media-only blank well as well as one empty pUCX (black bar) and three pUCXMG:azoR_Ec controls (grey bars), the latter of which were expected to be niclosamide-resistant but not metronidazole-sensitive as per Supplementary Figure S1). Data were derived from two biological repeats and error bars represent 1 S.D., while the black dashed lines indicate the cut-off that was imposed to define niclosamide resistance (A) or high-level sensitivity to metronidazole (B). The full screening dataset is available in Supplementary File S3; overall, 78% of niclosamide-challenged cultures achieved at least 50% culture turbidity (OD_600_) relative to control and 14% of metronidazole-challenged cultures were at least 80% growth-inhibited relative to control.

**Table 1:** Evaluation of the 21 pUCXMG:*eDNA* niclosamide-resistant and metronidazole-sensitive clones.

| NTR variant | NTR family | Metronidazole IC<sub>50</sub> (μM) | Metronidazole IC<sub>50</sub> (μM) +pRARE | In frame?<sup>a</sup> | GC content (%) | Number of rare codons<sup>b</sup> | Protein length (AA) |
| --- | --- | --- | --- | --- | --- | --- | --- |
| MhqN1 | MhqN | 17 ± 7 | 14 ± 5 | Yes | 63.9 | 4 | 205 |
| MhqN2 | MhqN | 23 ± 8 | 16 ± 5 | Yes | 64.1 | 7 | 205 |
| NfsB1 | NfsB | 92 ± 27 | 43 ± 28 | Yes | 63.3 | 6 | 216 |
| NfsB2 | NfsB | 217 ± 120 | 56 ± 21 | Yes | 62.5 | 7 | 282 |
| MhqN3 | MhqN | 92 ± 27 | 61 ± 26 | Yes | 67.0 | 4 | 205 |
| SagB1 | SagB | 189 ± 72 | 62 ± 15 | Yes | 66.5 | 11 | 187 |
| TdsD1 | TdsD | 130 ± 8 | 102 ± 33 | Yes | 67.8 | 5 | 201 |
| NfsA1 | NfsA | 105 ± 26 | 159 ± 38 | Yes | 67.2 | 14 | 260 |
| NfsB3 | NfsB | 767 ± 305 | 233 ± 91 | Yes | 55.3 | 9 | 216 |
| TdsD2 | TdsD | 873 ± 308 | 273 ± 46 | No | 44.6 | 10 | 192 |
| PnbA1 | PnbA | 618 ± 68 | 325 | Yes | 57.7 | 5 | 223 |
| TdsD3 | TdsD | 1200 ± 460 | 365 ± 83 | No | 42.7 | 9 | 192 |
| PnbA2 | PnbA | 1230 ± 260 | 555 ± 42 | No | 43.6 | 10 | 240 |
| PnbA3 | PnbA | 1110 ± 240 | 648 ± 14 | Yes | 39.7 | 13 | 220 |
| WrbA1<sup>c</sup> | WrbA | 2460 ± 500 | 1250 ± 430 | Yes | 64.6 | 2 | 194 |
| AzoR1<sup>c</sup> | AzoR | 2700 ± 980 | 1620 ± 740 | Yes | 63.8 | 5 | 208 |
| MhqN4 | MhqN | >5000 | 3890 ± 2760 | No | 45.9 | 12 | 233 |
| MhqN5 | MhqN | 295 ± 77 | ND<sup>d</sup> | Yes | 62.9 | 5 | 221 |
| NfsB4 | NfsB | 1300 ± 1100 | ND<sup>d</sup> | Yes | 70.8 | 5 | 216 |
| TdsD4 | TdsD | 1870 ± 730 | ND<sup>d</sup> | Yes | 53.7 | 8 | 192 |
| AzoR2<sup>c</sup> | AzoR | 2500 ± 1290 | ND<sup>d</sup> | Yes | 64.6 | 7 | 209 |
| **Controls** |  |  |  |  |  |  |  |
| pUCX: NfsA_Ec | NfsA | 73 ± 10 | 258 ± 109 | N/A | 52.6 | 2 | 240 |
| pUCX: NfsB_Ec | NfsB | 415 ± 91 | 284 ± 22 | N/A | 51.5 | 1 | 217 |
| Empty pUCXMG | N/A | >5000 | >5000 | N/A | N/A | N/A | N/A |

<sup>a</sup> Whether or not the *NcoI* site of pUCXMG comprised the likely start codon for the identified nitroreductase (NTR) variant.
<sup>b</sup> Incidence of six codons (AGG, AGA, AUA, CUA, CCC, GGA) underrepresented in *E. coli*, and for which matching tRNA genes are present on pRARE.
<sup>c</sup> Enzymes that are not part of the structurally-conserved nitroreductase superfamily, as defined by Akiva et al (2017).
<sup>d</sup> Insufficient growth in culture to enable IC<sub>50</sub> determination.
The apricot shading indicates enzymes from families that do not contain any previously-characterised bacterial nitroreductases, while the blue shading indicates enzymes from families that do not contain any previously-characterised nitroimidazole reductases.

### Protein production may be further enhanced by co-expression of rare tRNAs

SDS-PAGE analysis of each strain alongside a control expressing *E. coli nfsA* from plasmid pUCX revealed that the eDNA-derived nitroreductase levels were rather variable, with no over-expressed band being visible in some cases (**Figure 4A**). Codon analysis of the recovered sequences revealed that nearly all the recovered genes contained a higher number of rare *E. coli* codons than the native *nfsA* or *nfsB* genes (**Table** 1), which suggested that sub-optimal codon use might be impairing translation and thereby limiting their perceived activity in this host. To alleviate this issue, we co­transformed these strains with pRARE (a plasmid derived from the ROSETTA strain that supplements *E. coli* cells with rare tRNAs; Kirienko et al, 2004) and evaluated its effect on levels of enzyme expression (**Figure 4B**) and metronidazole IC_50_ (**Table** 1). Improvements in each parameter were observed for the majority of variants, with the sensitivity to metronidazole of *E. coli* cells bearing five different nitroreductases (NfsB2, NfsB3, SagB1, TdsD2, and TdsD3) enhanced by over 3-fold (**Table** 1). However, strains expressing four nitroreductases did not tolerate pRARE co-expression and grew poorly (AzoR2, MhqN5, NfsB4) or not at all (TdsD4), and pRARE also surprisingly impaired the metronidazole IC_50_ of the control strain bearing pUCX:nƒsA_Ec by over 3-fold (**Figure 4B**, **Table 1**). Overall, though, addition of pRARE was generally beneficial to the expression and activity of recovered nitroreductases, suggesting that steps to mitigate codon bias may add value to screening pipelines.

**Figure 4:**
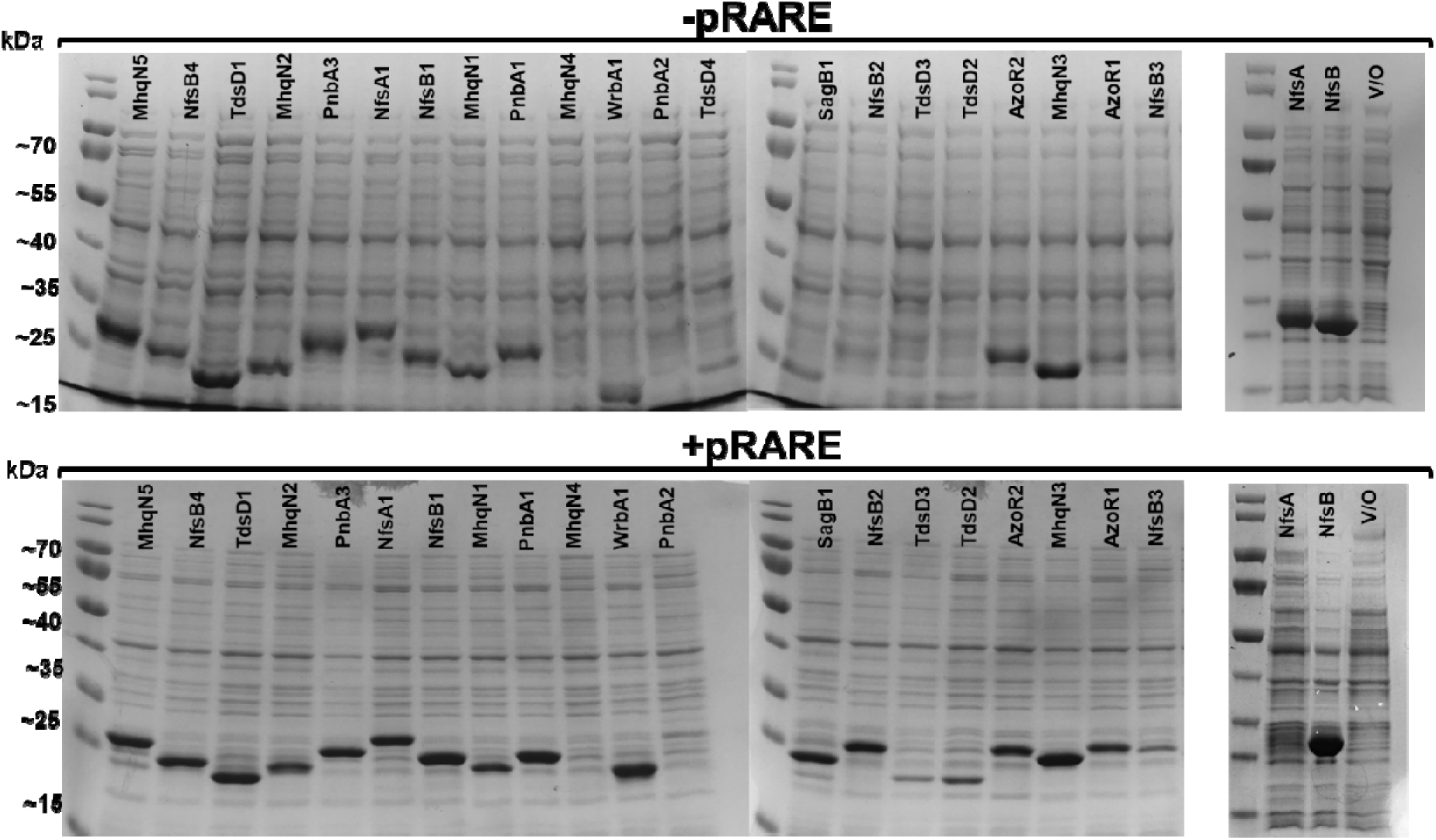
SDS-PAGE analysis of *E. coli* 7TL cells expressing captured nitroreductases. Enzymes were expressed from pUCXMG, without (top) or with (bottom) co-transformation by pRARE. Protein expression was induced and cultures incubated for 4.5 h, then cell densities normalised and loaded in the same order on each gel (except that there was no TdsD4 sample on the “+pRARE” gel as growth of the corresponding strain could not be achieved in liquid media). Control cultures of cells ± pRARE and expressing NfsA or NfsB from pUCX, or transformed by empty pUCX (V/O; vector only) were treated in identical fashion and analysed on a separate gel (rightmost panels).

### An embedded His_6_-tag allows purification of captured proteins without re-cloning

Based on the metronidazole IC_50_ data for the pRARE-containing strains (**Table** 1), we identified seven nitroreductases (MhqN1, MhqN2, NfsB1, NfsB2, MhqN3, SagBl and TdsDl) that conferred at least a four-fold greater sensitivity to metronidazole than observed for *E. coli* 7TL cells transformed with pUCX.nƒsB. All seven of the corresponding genes were ligated in frame with their start codons positioned within the Ncol/Fatl fusion site of pUCXMG, enabling us to test the utility of the embedded His_6_ tag for protein purification. In all cases, proteins were successfully purified when expressed from the pUCXMG screening plasmid (**Figure 5**), avoiding any need to re-clone the corresponding gene inserts into a specialised expression vector prior to protein purification. We noted there was a strong propensity for all proteins other than SagB1 to maintain a dimeric conformation even after boiling in SDS-PAGE loading buffer (**Figure 5**). We were also surprised to observe that TdsDl, MhqNl and SagBl were exclusively present in the insoluble fraction of lysates derived from the original 7TL screening strain, and it was instead necessary to transfer the corresponding pUCXMG plasmids to the specialised *E. coli* expression strain BL21 to achieve soluble protein preparations. This additional transfer step could presumably have been avoided by conducting our metagenome screening in a BL21-derived host strain.

**Figure 5:**
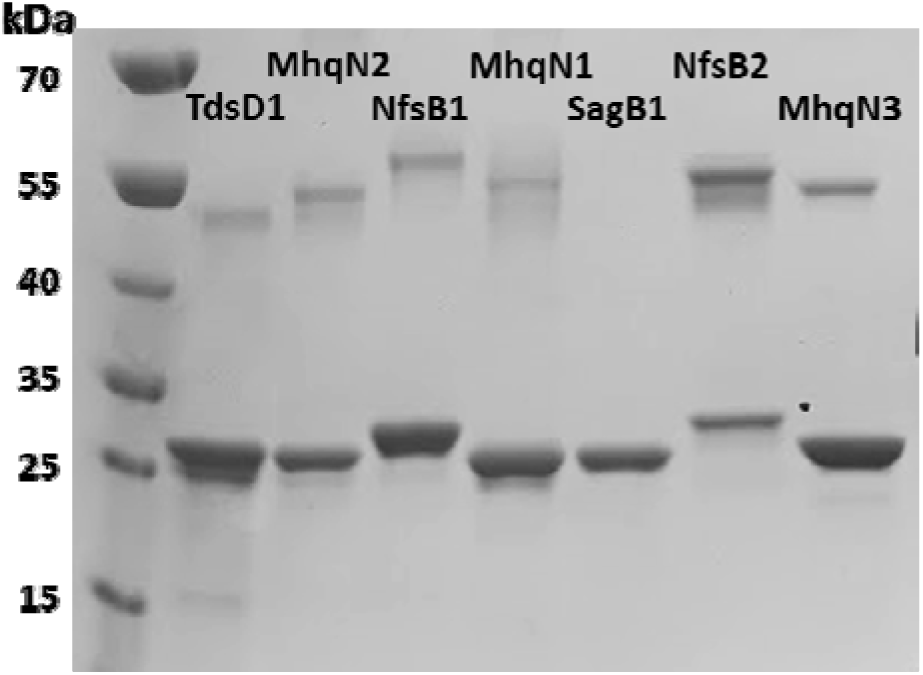
SDS-PAGE of recombinant His_6_-tagged nitroreductases. Each nitroreductase was purified by standard Ni/NTA chromatography post-expression from each respective pUCXMG cloning plasmid. Five micrograms of purified protein were loaded per lane.

### The top metagenome-derived nitroreductase MhqN2 outperforms the canonical nitroreductase E. coli NfsB for targeted cell ablation in zebrafish

We have previously observed that certain bacterial nitroreductases express poorly, or not at all, in eukaryotic models, which we attribute to the potential for nitroreductase substrate promiscuity to disrupt primary metabolic pathways (Sharrock et al, 2022). To determine whether any of our top seven eDNA-derived nitroreductases could be used for targeted cell ablation in zebrafish, we attempted to create transgenic zebrafish lines co-expressing each enzyme with a YFP reporter. For this, transgenic UAS reporter/effector lines, *Tg(5xUAS:YFP-2A-nitroreductase,he:tagBFP2)* fish were generated as previously described (Sharrock et al., 2022). Each UAS line was crossed to a previously established Gal4 enhancer trap driver line, *Et(2xNRSE-Mmu.fos:KALTA4)gmc617Et* (Xie et al., 2012), to restrict nitroreductase and YFP co-expression to the same set of targeted neurons. The transgenic lines that were recovered for TdsDl, NfsBl or NfsB2 did not express YFP at detectable levels and were not further investigated. However, we successfully generated distinct transgenic zebrafish lines co-expressing the YFP reporter and MhqN1, MhqN2, MhqN3 or SagB1.

To assay the abilities of these nitroreductase variants to induce cell ablation, larvae from each strain were subjected to a titration of metronidazole concentrations (0,1, 5 or 10 mM) at 5 days post­fertilisation (5 dpf). After 48 h of exposure, residual levels of YFP expression in 7 dpf larvae were quantified using a TECAN fluorescence microplate reader as previously described (Sharrock et al., 2022). Partial ablation was apparent for the lines expressing MhqNl and SagBl, and near complete ablation for the line expressing MhqN2 at all concentrations tested (p<O.OOOl relative to control) (**Figure 6**). The MhqN2 line was then subjected to a further titration of metronidazole concentrations (0.1, 0.2 and 0.5 mM; **Figure 6E**) that enabled calculation of an absolute EC_50_ of 430 µM. This was ∼5- fold more effective than a previously generated control line co-expressing the benchmark nitroreductase *E. coli* NfsB and mCherry, *Tg(UAS:NTR-mCherry)c264* (Davison et al., 2007) in the same neuronal target cells (i.e., crossed to the same Gal4 driver, *gmc617Et;* Xie et al, 2012), which yielded an absolute EC_50_ of 2.3 mM metronidazole (**Figure 6D**).

**Figure 6:**
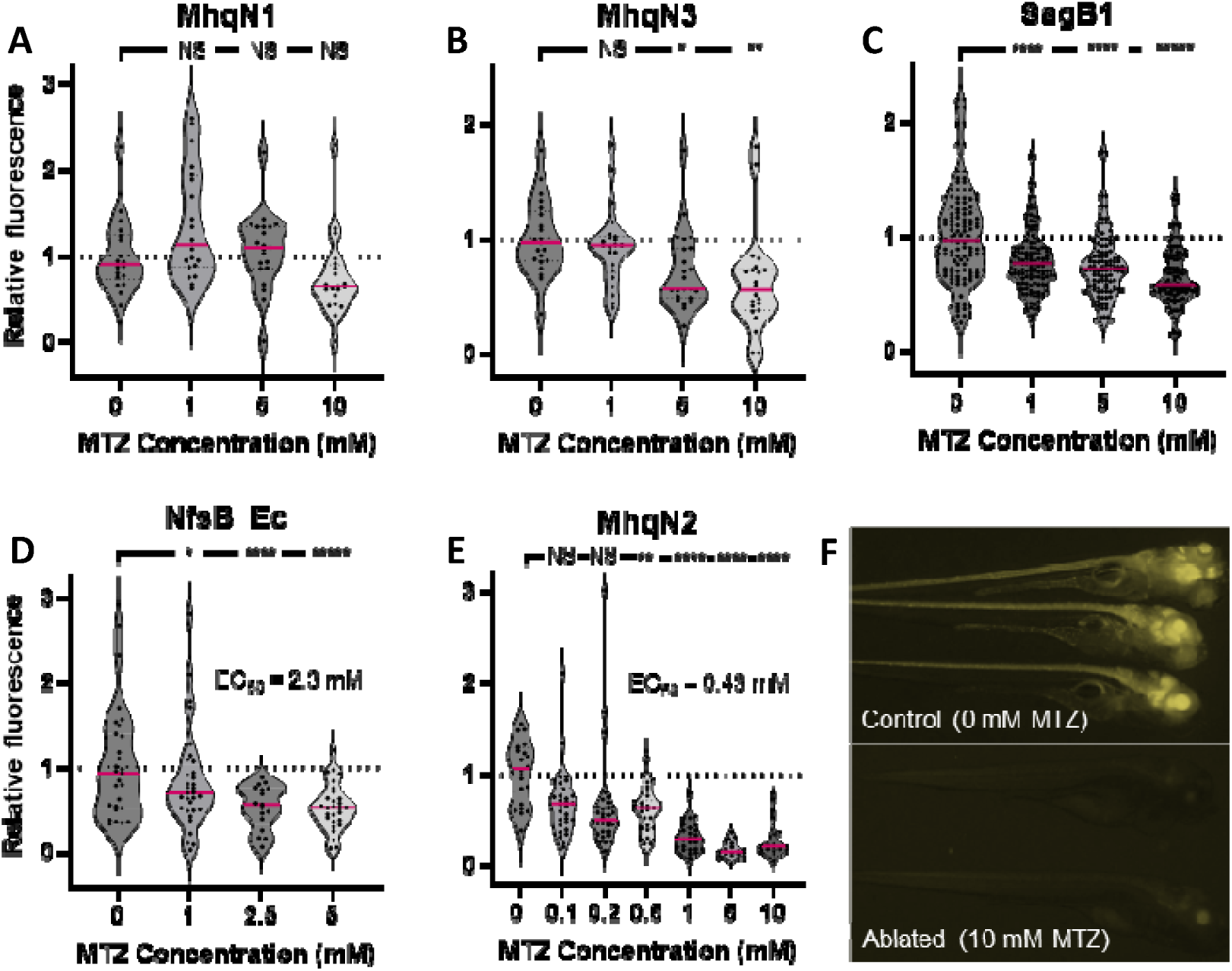
Cell ablation efficacy in transgenic zebrafish for neuronal cells expressing lead nitroreductase candidates. **A-E)** Transgenic zebrafish larvae co-expressing the indicated nitroreductase and either yellow fluorescent protein **(A-C,E)** or mCherry **(D)** in cells of the central nervous system were exposed to a range of metronidazole concentrations to assess relative cell ablation efficacy. In initial tests, the MhqN2 line **(E)** showed >50% ablation at 1 mM metronidazole and was exposed to lower concentrations to enable measurement of an absolute EC_50_. Bonferroni-corrected p’-values relative to the control condition (0 mM metronidazole) are indicated by asterisks: *p<0.05, **p<0.01, ***p<0.001, ****p<0.0001 (NS = not significant). **F)** Micrographs of MhqN2 expressing zebrafish larvae after 48 hours of exposure to control media (above) or media containing 10 mM metronidazole (below).

## Discussion

*We* describe here a broadly-applicable strategy to generate small-insert eDNA libraries that are greatly enriched for genes with their start codons placed an optimal distance downstream of a strong *E. coli* promoter and ribosome binding sequence. This enables efficient selection or screening for weak phenotypes that require high levels of gene expression to manifest, as per the niclosamide and metronidazole converting nitroreductases exemplified here. Although a small proportion of our recovered nitroreductase genes initiated from internal start codons rather than at the *Nco\-Fat\* ligation point, suggesting they might have been recoverable from standard eDNA libraries, over 80% of selected genes were ligated in-frame at the *Ncol-Fatl* fusion point, consistent with the majority having required the boosted expression our cloning strategy provides. We anticipate that this boosted expression will provide substantial benefit to enzyme discovery campaigns that employ ultra-high throughput fluorescence activated cell or droplet sorting technologies, as these impose an extreme requirement for strong signals from very small reaction volumes (Sheludko and Fessner, 2020). However, by minimising the incidence of non-expressing inserts our approach will also benefit screens that only have low to moderate throughput and hence require a high ‘hit’ frequency (Ferrer et al, 2016) (e.g., discovery of substrate-converting enzymes using thin-layer chromatography; Rabausch et al, 2013). We also showed that a N-terminal Hisg-tag could be embedded in the vector to streamline purification and biochemical evaluation of recovered enzymes. While it is possible that some desirable enzyme variants may not tolerate a purification tag in this position, a pragmatic consideration is that screening with a tag in place will select for enzymes that are more likely to be amenable to biochemical characterisation.

Our strategy was exemplified using eDNA from soil, which can represent many thousands of bacterial species per gram (Roesch et al, 2007; Crits-Christoph et al, 2018), but we anticipate it will be readily applicable to other sources, e.g. to interrogate the human gut microbiome to detect drug­modifying enzymes, or to identify enzymes with bioremediation potential from polluted environments. We believe it will also hold great value in discovering individual enzymatic tools for synthetic biology, while similarly-designed libraries that employ larger insert sizes may also prove useful for capturing entire operons, e.g. for discovery of natural product gene clusters by screening for characteristic ‘beacon’ genes (Baltz, 2017). However, while our approach may offer substantial advantages in activating the expression of operons that might otherwise be silent (Rutledge and Challis, 2015; Mao et al, 2018), it will not preferentially clone complete operons over partial ones, so it must be considered that small operons are far more likely to be recovered intact than large ones.

In analysing the occurrence of (C)ATG start codons in genome-sequenced bacteria, we made a surprising observation that cytosine is over-represented at the -1 position relative to ATG start codons. The heightened frequency of cytosines in this position cannot be attributed solely to the GC content of the host organism, as (C)ATG start codons appear nearly twice as frequently as (G)ATG codons. We consider it plausible that the higher incidence of palindromic CATG sequences could reflect secondary structures that may form around the translational start point, with possible regulatory roles (e.g., it was recently shown that reducing mRNA secondary structure around the start codon substantially increased expression of the fluorescent reporter mNeonGreen in both *Saccharomyces cerevisiae* and *E. coli;* Hector et al, 2021). Irrespective, while this is a beneficial phenomenon for our *FatI* cloning strategy in terms of increased likelihood of capturing start codons, it does reflect that our strategy is likely to be biased toward capture of genes from GC-rich bacteria. The lower GC content of *E. coli* (50.8%; Blattner et al, 1997) relative to the majority of recovered nitroreductases likely contributed to the incidence of rare codons and poor expression of some of these enzymes. We showed that expression of some nitroreductases was improved by co­transformation of the host with pRARE; but in some other cases this actually diminished nitroreductase activity. Thus, for groups seeking to maximise gene recovery, there may be value in conducting parallel screens of a host strain transformed by the eDNA library alone, alongside another host that has been co-transformed with pRARE. It is possible that addition of other genes that facilitate heterologous expression, e.g. increase chaperone production, might also improve the recovery of genes from distant phyla.

The great strength of conducting functional metagenomic screens or selections is that one is not limited to only the ‘known unknowns’, i.e. homologues of proteins already known to possess the activity of interest. The power to recover novel biocatalysts was on display here. As far as we are aware, this is the first experimental demonstration of any nitroreductase activity for bacterial enzymes from the unrelated SagB (azole biosynthesis) and WrbA (quinone oxidoreductase) families; and while there has been one report apiece of nitroreductase activity from TdsD (Takahashi et al, 2009) and MhqN (Takeda et al, 2007) family members (which share a conserved fold with the better-known NfsA, NfsB and PnbA nitroreductases; Akiva et al, 2017), no activity had previously been described with nitroimidazole substrates. Despite this, our two most active metronidazole reductases in an *E. coli* host (MhqN1, MhqN2) were both from the MhqN enzyme family. The value in recovering a broad range of metronidazole reductases was evident when testing in a transgenic zebrafish model of cellular ablation (an environment where we have previously observed only a subset of otherwise-promising nitroreductases to function; Sharrock et al, 2022). In this model, MhqN2 appeared ∼5-fold more effective than the canonical nitroreductase, *E. coli* NfsB, which was previously found to be insufficiently active for ablation of certain cell types, e.g., dopaminergic neurons (Godoy et al, 2015), cone photoreceptors (Fraser et al, 2013), and macrophages (Sharrock et al, 2022). Importantly, that MhqN2 enables effective ablation at non-toxic Mtz dosages (<1mM; Sharrock et al., 2022) opens opportunities for chronic ablation paradigms, i.e. continuous metronidazole exposure for inducible modelling of long-term degenerative diseases.

Not only does accessing a far greater breadth of diversity increase the chances of uncovering an enzyme that is substantially better than any native enzymes previously known (as was the case here), it also provides a broad range of starting points for directed evolution to further improve the desired activity. This breadth will be beneficial to avoid local maxima that have potential to stall directed evolution campaigns when the initial levels of diversity are low (Gupta and Tawfik, 2008; Packer and Liu, 2015). Moreover, functional metagenomics and directed evolution both usually require efficient high-throughput screens or selections, and the same basic pipeline can often be applied to further enhance activity by evolving the top enzymes recovered from eDNA library screening. One key difference is that directed evolution usually seeks to discriminate between closely related variants, whereas metagenome screening can uncover entirely unrelated enzymes, and hence is more likely to encounter substantial discrepancies in relative expression levels. The strategies we describe here to boost expression of genes captured in eDNA libraries should mitigate this impact and thereby facilitate combined discovery and evolution campaigns.

### Significance

Modern DNA sequencing technologies are probing deeper than ever before into the ‘microbial dark matter’ within complex environments such as soil. However, biochemical characterisation of the diversity of proteins encoded by this sequence is lagging far behind. Functional screening of environmental DNA is an attractive strategy to discover new enzymatic activities without requiring preconceptions of the types of enzymes likely to be catalysing the desired chemistry, which will perforce be heavily biased toward previously-characterised protein families. We describe here an environmental DNA cloning strategy that ensures potential start codons are placed an optimal distance downstream of a strong host-appropriate promoter and ribosome binding sequence, and show that it greatly enriches for captured genes that express efficiently in the new host cell. This overcomes an important and long-established roadblock to effective functional screening. In particular, it provides access to weak promiscuous activities that require high-level gene expression to confer a detectable phenotype, and yet might hold particular value for biotechnology. We exemplify that here by recovering 21 enzymes from eight different families that are each active with non-biological molecules, able to detoxify the antibiotic niclosamide and activate the prodrug metronidazole. This collection included enzymes from two families that had not previously been implicated in bacterial nitro-reduction. Our best-performing enzyme in a transgenic zebrafish model of targeted cellular ablation was effective at ∼5-fold lower concentrations of metronidazole than the previous benchmark enzyme *E. coli* NfsB, illustrating the power of an unbiased screening approach to recover desirable activities.

## STAR Methods

### Strains, media, chemicals and plasmids

All bacterial screening and growth assays were performed using *E. coli* 7NT or its *λ*DE3 lysogenised derivative 7TL as described in this study. 7NT was derived from the standard laboratory strain W3110 by scarless in-frame deletion of seven candidate nitroreductase genes *(nfsA, nfsB, azoR, nemA, yieF, ycaK* and *mdaB)* and the efflux pump gene *tolC* (Copp et al., 2014). As detailed below, the FatI library was initially prepared in *E. coli* DHB10 and proteins were purified from either *E. coli* BL21 or 7TL. All chemicals were sourced from Duchefa Biochemie unless otherwise stated. Bacterial cultures were grown and assessed in Lysogeny Broth (LB) amended with antibiotics as appropriate for plasmid maintenance (100 µg.mL^-1^ ampicillin for pUCX or pRSETB, 20 µg.mL^-1^ gentamycin for pUCXMG, and/or 30 µg.mL^-1^ chloramphenicol for pRARE). Plasmid pUCX was previously generated in house (Prosser et al, 2013; Addgene plasmid #60681), pRSETB bearing a soil eDNA library was kindly provided by Nadia Parachin and Marie Gorwa-Grauslund (Parachin and Gorwa-Grauslund, 2011), pRARE was as described by Kirienko et al (2004), and pUCXMG was created for this study as described below.

### Bioinformatics

All scripts developed for bioinformatic analyses are available at the following GitHub link: https://github.com/michhrich/metagenomic-library-rich-et-al-2023. For this work, 21,675 complete assembled bacterial genomes were downloaded from NCBI (https://www.ncbi.nlm.nih.gOv/genome/browse#!/prokaryotes/) in June 2021. A Python script (https://github.com/michhrich/metagenomic-library-rich-et-al-2023/blob/main/supplementary-script-sl) was then used to extract and compile elements from individual chromosomes within each genome (by excluding records with ‘plasmid in the record description, including the position -1 to 3 sequences at the start of each predicted open reading frame for each annotated CDS. For genomes containing multiple chromosomes, the results were combined into a single record using a second Python script (https://github.com/michhrich/metagenomic-library-rich-et-al-2023/blob/main/supplementary-script-sl). The scripts are additionally available as Supplementary File S1, and their combined output was compiled into an Excel worksheet and provided as Supplementary File S2.

### E. coli growth inhibition assays and IC_50_ analysis

Day cultures were established by adding 150 µl of overnight culture to 3 ml fresh LB amended with ampicillin and 50 µM IPTG in a 15 ml tube, for each strain to be assessed. Day cultures were incubated to induce protein expression for 2 h at 30 °C with shaking at 200 rpm. For growth inhibition assays, 40 µl aliquots of culture were added to individual wells of a 384 well plate containing 40 µl LB, either unamended as an unchallenged control, or amended with metronidazole or niclosamide at twice the desired final concentration. Culture turbidity (OD_600_) was read initially (T_o_), and again following 4 h incubation at 30 °C, 200 rpm (T_4_). Percentage growth for challenged strains was then calculated by subtracting the T_0_ value from the T_4_ for each well, converting any negative values to zero, then dividing the data for challenged wells by the corresponding data for the unchallenged control. For IC_50_ assays, a range of growth inhibition data were calculated in an equivalent fashion, from replicate cultures across a two-fold dilution series of 800 µM to 24 nM metronidazole, or 50 µM to 1.5 nM niclosamide. Final IC_50_ values were calculated from three biological replicates each comprising two technical replicates using Graphpad Prism software.

### Screening of pRSETB soil eDNA library

Initial screening of the pRSETB soil eDNA library created by Gorwa and Grauslund (2011) was performed to >3-fold coverage in *E. coli* 7TL cells on LB agar amended with ampicillin, 0.5 µM niclosamide and 50 µM IPTG, and yielded three different eDNA inserts containing *tolC*-like genes. The library was subsequently rescreened on LB agar amended with ampicillin, 50 µM IPTG, niclosamide and the TolC inhibitor phenylalanine-arginine beta naphthylamide (PAβN; Lomovskaya et al, 2001), with or without addition of 1 mM MgSO_4_ to mitigate the membrane permeabilising effects of PAβN (Lamers et al, 2013). Two paired concentrations of PAβN and niclosamide were used, 100 µM PAβN and 0.1 µM niclosamide, or 50 µM PAβN and 0.2 µM niclosamide (each empirically found to prevent the growth of *E. coli* 7TL cells expressing *tolC*-like genes recovered in the initial screen, but permit the growth of 7TL transformed by pUCX bearing *E. coli nfsB)*.

### Design and assembly of plasmid pUCXMG

The pUCXMG vector was generated from the pUCX vector backbone with a replacement of the ampicillin resistance marker by a gentamycin resistance cassette and introduction of a modified multiple-cloning site, containing a Hise tag and a downstream NcoI restriction site. For replacement of the antibiotic resistance gene, the pUCX vector was amplified with forward primer CTGTCAGACCAAGTTTACTCATATATACTTTAGATTGATTTAAAAC and reverse primer ACTCTTCCTTTTTCAATATTATTGAAGC and assembled with a synthetic gentamycin cassette ordered from Twist Bioscience containing 5’ and 3’ 20-bp pUCX homology arms, using NEBuilder® HiFi DNA Assembly (New England Biolabs). A synthetic cloning site comprising an NcoI recognition sequence flanked by XbaI and HindIII restriction sites was ordered from Twist Bioscience and used to replace the XbaI-HindIII region of the pUCX multiple cloning site by restriction cloning. The complete sequence of the final pUCXMG plasmid is available in **Supplementary Figure S2**.

### Generation of a high-expression soil eDNA library using Fatl partial digestion

Metagenomic DNA was extracted from soil collected from a private residence in Holloway Road, Wellington, New Zealand as per the protocol of Stevenson et al (2022). The eDNA was further purified to remove humic inhibitors by electrophoresis through an agarose gel (1% w/v low-gelling-temperature agarose (Sigma Type VII) in 1× TAE buffer) for 1 h at 150 V, 4 °C. The agarose gel was post-stained with SYBR Safe DNA stain (Thermo Fisher), and the high-molecular-weight DNA was sliced from the gel and digested with β-Agarase I (New England Biolabs) for 1 h at 42 °C. The eDNA was then purified from the digested solution by precipitating with 60% isopropanol + 300 mM sodium acetate pH 5.2 in microcentrifuge tubes. Tubes were centrifuged at 17,000 *g,* supernatants discarded, and pellets washed with -20 °C 70% EtOH (v/v). After this, supernatants were discarded and pellets air dried for 5 min, then resuspended in 10 mM Tris-HCl pH 8.0 and DNA concentrations determined using a nanodrop spectrophotometer.

For library generation, eDNA was partially digested by adding 1.1 U FatI/µg eDNA and incubating at 55 °C until test reactions revealed a substantial ‘smear’ in the 0.5 to 5 kb range when visualised on a 1% agarose gel. The digested eDNA was electrophoresed on a low melting temperature agarose gel with a sacrificial sample in the lane next to the markers being stained for visualisation and the *ca.* 0.6-1.4 kb range marked. The neighbouring (unstained) lanes were then aligned against the marks and the equivalent regions excised, then DNA fragments extracted and purified as described above. The extracted eDNA fragments were then ligated with pUCXMG vector that had been linearised by NcoI digestion in a 2:1 ratio with overnight at 4 °C. The ligated DNA was co-precipitated with yeast tRNA (1 µl of 1 µg/µl tRNA per 5 µl ligation mixture) using isopropanol/sodium acetate followed by a 70% ethanol wash and resuspension in 10 mM Tris-HCl pH 8.0 as above. The resulting FatI library ligation was used to transform electrocompetent *E. coli* DH10B cells that were then plated onto LB agar amended with gentamycin; a serial dilution of small aliquots on 90 mm plates to estimate library size, and the remainder on a 150 mm plate. Cells were collected from the latter by adding 2 ml LB broth, scraping, and transferring the liquid to a centrifuge tube. Centrifugation was performed for 1 h at 2,400 *g,* after which the supernatant was discarded and the pellet resuspended in fresh LB broth to form a thick slurry. Aliquots from the slurry were miniprepped to provide a DNA level library and the remainder mixed 1:1 with 80% glycerol (v/v) and snap frozen at -80 °C as a renewable stock. Insert rates were estimated by colony PCR using 56 colonies randomly selected from the serial dilution plates used to estimate library size, with the primers pUCX_for (GACATCATAACGGTTCTG) and pUCX_rev (GTTTCACTTCTGAGTTCG) that flank the NcoI cloning site of pUCXMG.

### Selection and evaluation of nitroreductases from the Fatl eDNA library

*E. coli* 7TL cells transformed with the FatI eDNA library were plated on LB agar amended with gentamycin, 0.5 µM niclosamide, and either 5 or 50 µM IPTG. Any resulting colonies were individually picked into fresh LB amended with gentamycin in 96 well microplates and the resulting cultures subjected to niclosamide and metronidazole growth inhibition assays as described above. Niclosamide-resistant and metronidazole-sensitive clones were miniprepped and Sanger sequenced by Macrogen (South Korea) in both orientations using primers pUCX_for and pUCX_rev (details above). Sequenced inserts were analysed against the NCBI non-redundant protein sequence database using BLASTx, in each case revealing a predicted protein sequence annotated as a nitroreductase or NAD(P)H-dependent oxidoreductase. These were assigned to a nitroreductase sub-family (Akiva et al., 2017) by BLAST search against the Structure-Function Linkage Database (http://sfld.rbvi.ucsf.edu/archive/django/index.html; now archived) or else annotated as members of the non-homologous AzoR or WrbA enzyme families.

### Protein purification and SDS-PAGE analysis

Hisētagged proteins were purified using Ni/NTA columns (Novagen), following expression in the *E. coli* 7TL screening strain (or *E. coli* BL21 for TdsDl, MhqNl or SagBl). Inocula from overnight cultures were incubated in 50 ml of fresh LB containing gentamycin at 37 °C with shaking at 200 rpm until a turbidity of OD_600_ of 0.5 was achieved. Cultures were then chilled on ice for 15 min, IPTG added to a final concentration of 0.5 mM, and then incubated at 18 °C for 16 h. Following centrifugation, pellets were resuspended in 20 ml HisBind buffer (Novagen) and cells lysed by French pressing, with supernatants from a further centrifugation step being applied to the Ni/NTA columns. Post purification, purity was assessed by SDS-PAGE using 12.5% acrylamide gels with 5 µg protein loaded per lane and bands visualised by staining with Coomassie Brilliant Blue.

For SDS-PAGE analysis of cells expressing nitroreductase genes, cultures of each strain were established as above then incubated for 4.5 h at 30 °C post-addition of 50 µM IPTG, after which cells were spun down, resuspended in 50 µl LB and normalised to an OD_600_ of 5. A 20 µl volume of each cell resuspension was boiled in SDS-PAGE loading buffer and loaded per lane and bands were visualised by staining with Coomassie Brilliant Blue.

### Evaluation of lead nitroreductases in transgenic zebrafish

A subset of nitroreductases shown to effectively convert metronidazole in bacteria were used to create novel zebrafish transgenic lines. UAS-based reporter/effector transgenes for co-expressing nitroreductase variants and the yellow fluorescent protein tagYFP were assembled and corresponding transgenic lines created as previously described (Sharrock et al., 2022). **Table 2** lists transgenic details for the Gal4 driver and UAS;nitroreductase lines utilized here. All UAS lines were crossed to the same previously established Gal4-based driver line, *Et(2xNRSE-Mmu.fos:KALTA4)gmc617* (Xie et al., 2012), in order to test cell ablation efficacy in the same population of neurons targeted by the *gmc617* line. Relative YFP expression levels were quantified following exposure to metronidazole at the indicated concentrations using an established fluorescence plate reader assay (Walker et al., 2012). Evaluations of all nitroreductase strains involved quantifying fluorescence before and after metronidazole exposure to allow normalization per each individual fish (i.e., with relative fluorescence expressed as the post-metronidazole fluorescence reading divided by the pre-metronidazole fluorescence reading). Evaluation of *E. coli* NfsB efficacy involved a post-metronidazole reading only, as previously described (Sharrock et al., 2022). All data was processed and plotted using GraphPad Prism. Absolute EC50 values - i.e., the concentration predicted to elicit 50% cell ablation - were calculated from dose-response data using an online EC50 calculator (https://www.aatbio.com/tools/ec50-calculator) and solving for y = 0.5. Multiple comparison corrected p-values were used for statistical comparisons. Micrographs demonstrating metronidazole-induced cell ablation efficacy in anesthetized zebrafish larvae were collected on a MVX10 Olympus fluorescence stereoscope with an Olympus DP72 camera (MhqN2), or an MV1000 Olympus confocal microscope, as previously described (F. *coli* NfsB; Ariga et al, 2010).

## Supporting information

Supplementary File S1 - Python Scripts

Supplementary File S2_evaluation of bacterial start codons

Supplementary File S3_collated growth inhibition data

## Acknowledgements

This work was supported by the Royal Society of New Zealand Marsden Fund (contract VUW1902; D.F.A., J.G.O.), the Health Research Council of New Zealand (contract 18-532; D.F.A.) and the US National Institutes of Health (R010D020376 and RF1MH126731 awards to J.S.M. and D.F.A., and a P30 core grant to the Wilmer Eye Institute, P30EY001765).

## Author contributions

**Conceptualisation:** MHR, JSM, DFA. **Methodology:** MHR, AVS, TSM, FM, ASB, HRLH, EMW, JNC, RFL, CNH, ĲS, JGO, MTS, JSM, DFA. **Software:** MHR. **Validation:** MHR, AVS, ASB, HRLH, EMW. **Formal Analysis:** MHR, AVS, TSM, FM, ASB, HRLH, JSM, DFA. **Investigation:** MHR, AVS, TSM, FM, ASB, HRLH, EMW, JNC, RFL, JJBF, CNH. **Resources:** US, JGO, MTS, JSM, DFA. **Data Curation:** MHR, MTS, JSM, DFA. **Writing-Original Draft:** MHR, DFA. **Writing - Review & Editing:** MHR, AVS, TSM, FM, ASB, HRLH, EMW, JNC, RFL, JJBF, CNH, US, JGO, MTS, JSM, DFA. **Visualisation:** MHR, AVS, FM, ASB, HRLH, JJBF, JSM, DFA. **Supervision:** JGO, MTS, JSM, DFA. **Project Administration:** DFA, JSM. **Funding Acquisition:** DFA, JGO, JSM.

## Declaration of Interests

The authors declare no competing interests.

## SUPPLEMENTARY MATERIALS

**Supplementary Figure S1.**
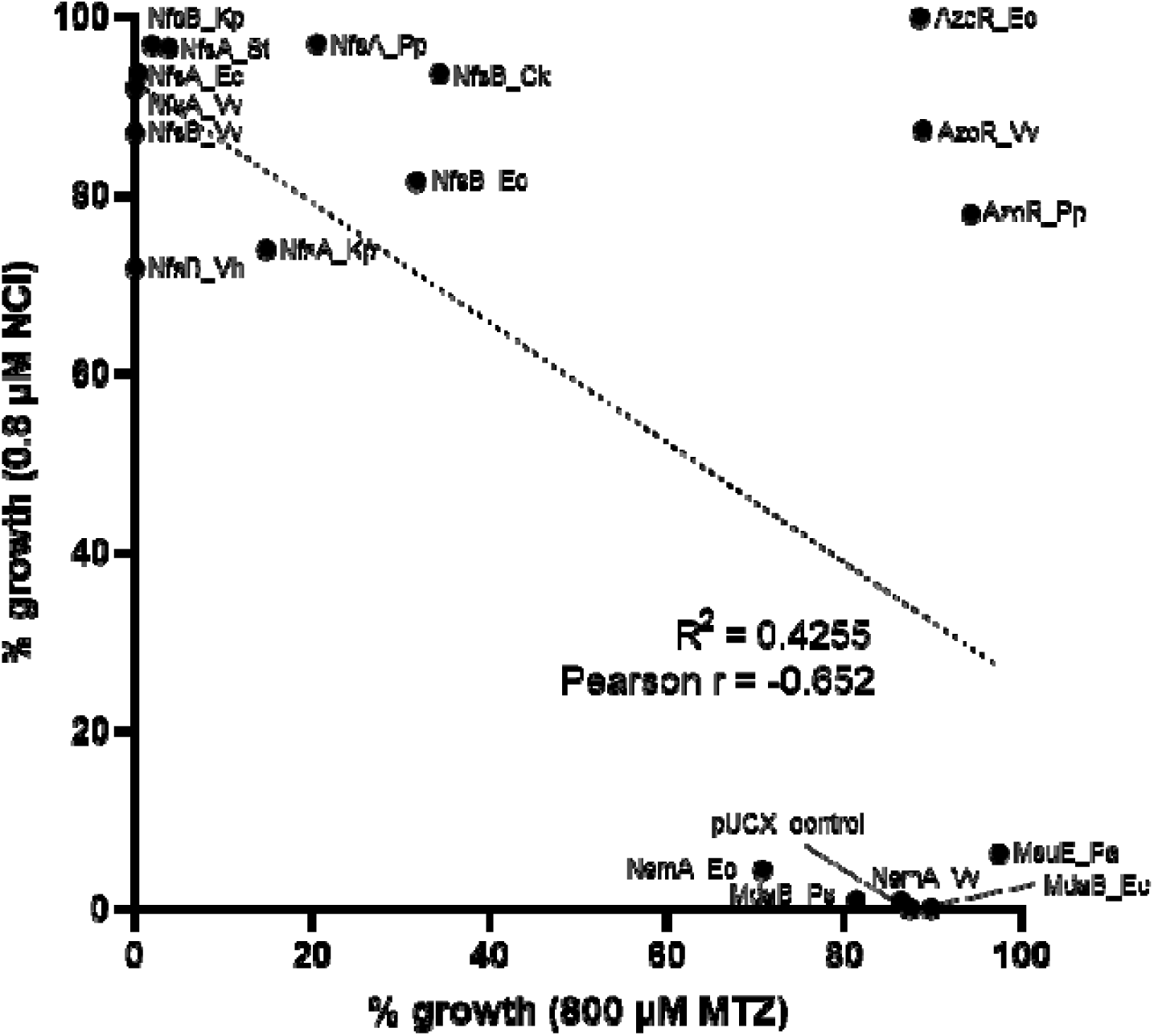
Comparison and correlation of niclosamide detoxification and metronidazole activation by *E. coli* 7NT strains expressing 18 nitroreductase candidates (from the NfsA, NfsB, AzoR, NemA, MsuE or MdaB enzyme families) from plasmid pUCX. Suffixes indicate the bacterial species each nitroreductase was derived from (Ck = *Citrobacter koseri*, Ec = *E. coli*, Kp = *Klebsiella pneumoniae*, Pa = *Pseudomonas aeruginosa*, Pp = *Pseudomonas putida*, Ps = *Pseudomonas syringae*, St = *Salmonella typhi*, Vh = *Vibrio harveyi*, Vv = *Vibrio vulnificus*; Copp et al, 2017).

**Supplementary Figure S2:**
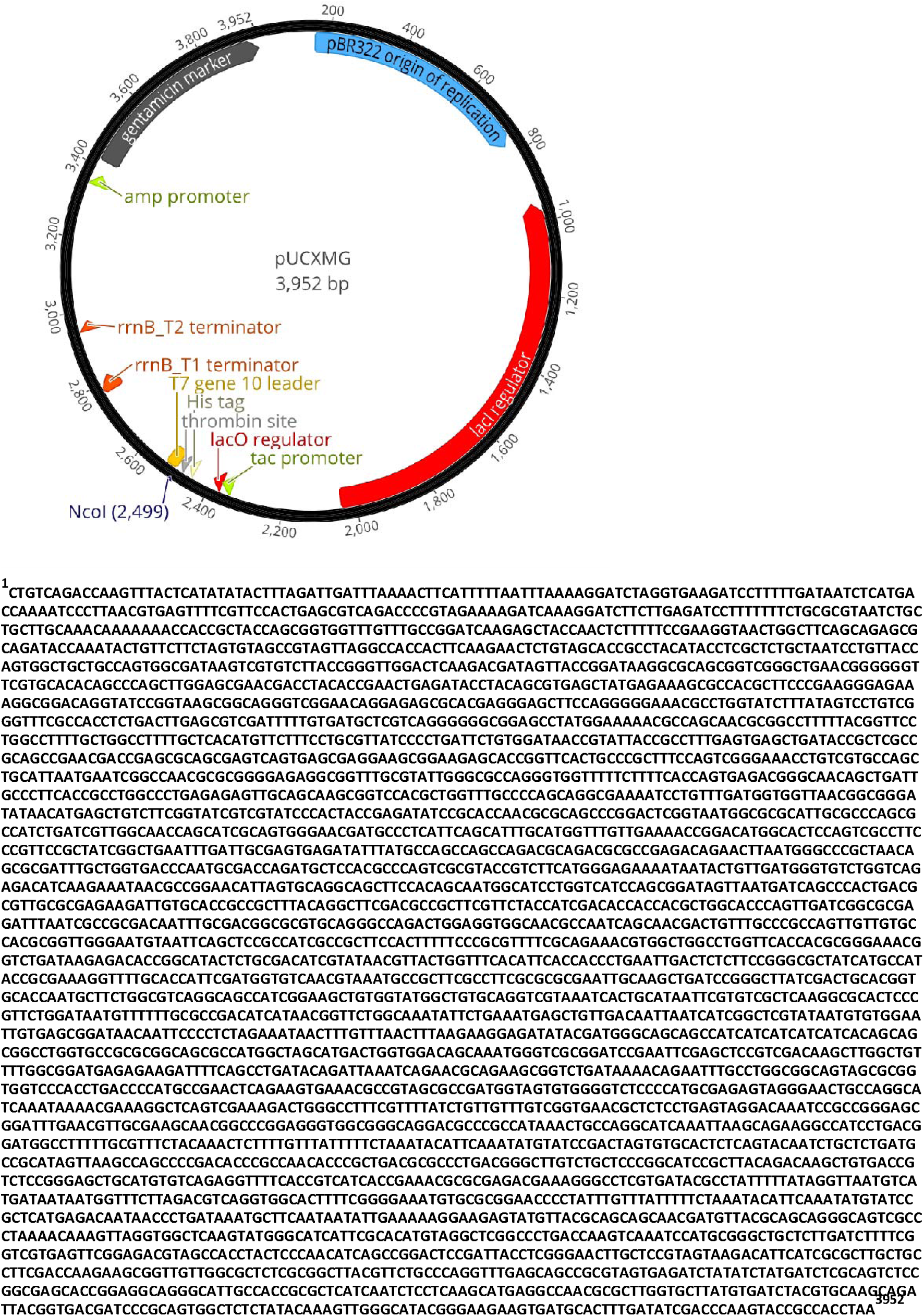
Map of plasmid pUCXMG. Salient features are highlighted using standard abbreviations, and the complete plasmid sequence is provided below the map.

**Supplementary Figure S3:**
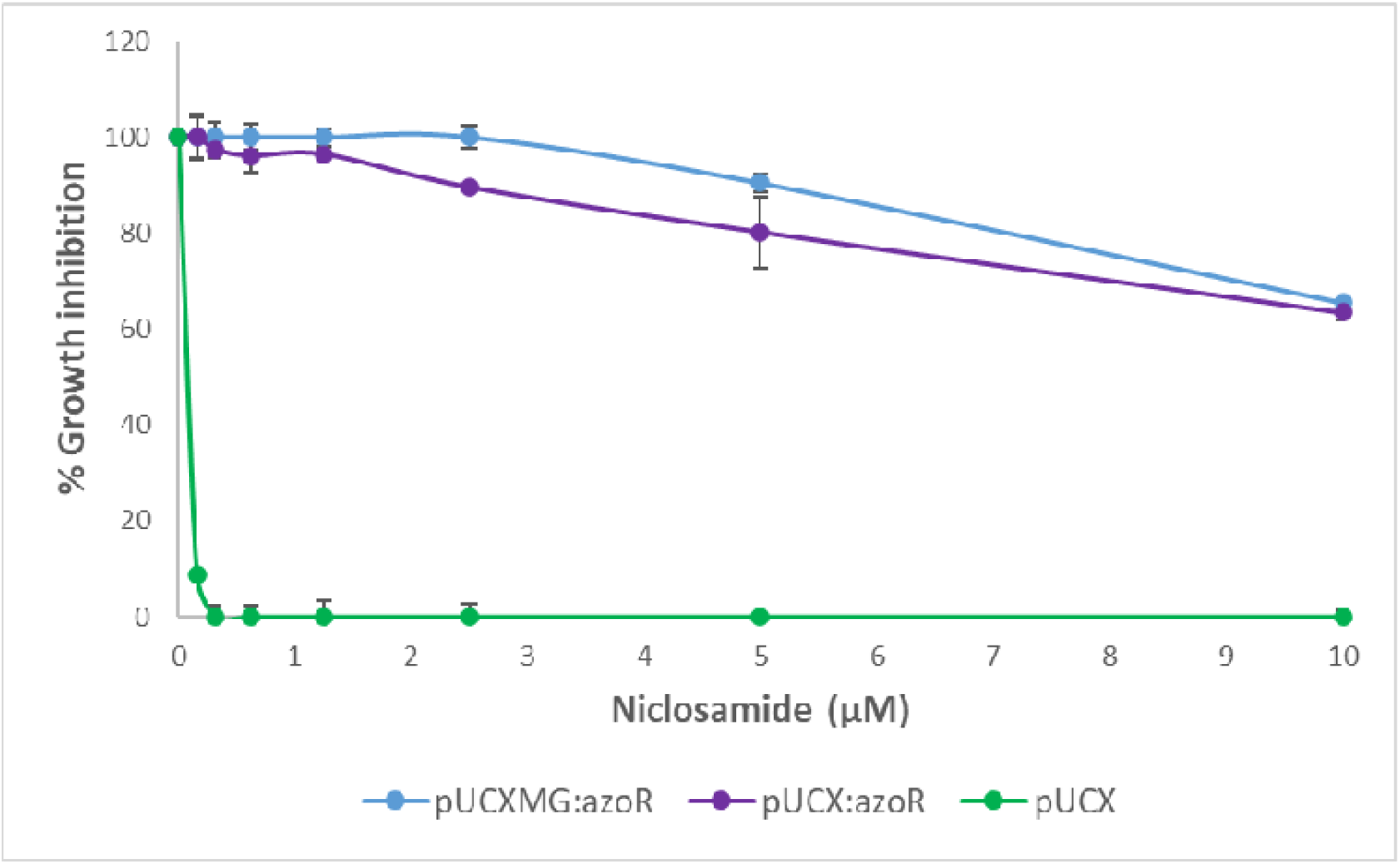
Replicate cultures of *E. coli* 7TL cells transformed by empty pUCX (green), pUCX expressing *azoR* (purple), or pUCXMG expressing *azoR* (blue) were induced with 50 µM IPTG and challenged with 0-10 µM niclosamide, with percentage growth (OD_600_) relative to the unchallenged control recorded after four hours. Data were derived from two technical replicates ± S.D.

**Supplementary Figure S4:**
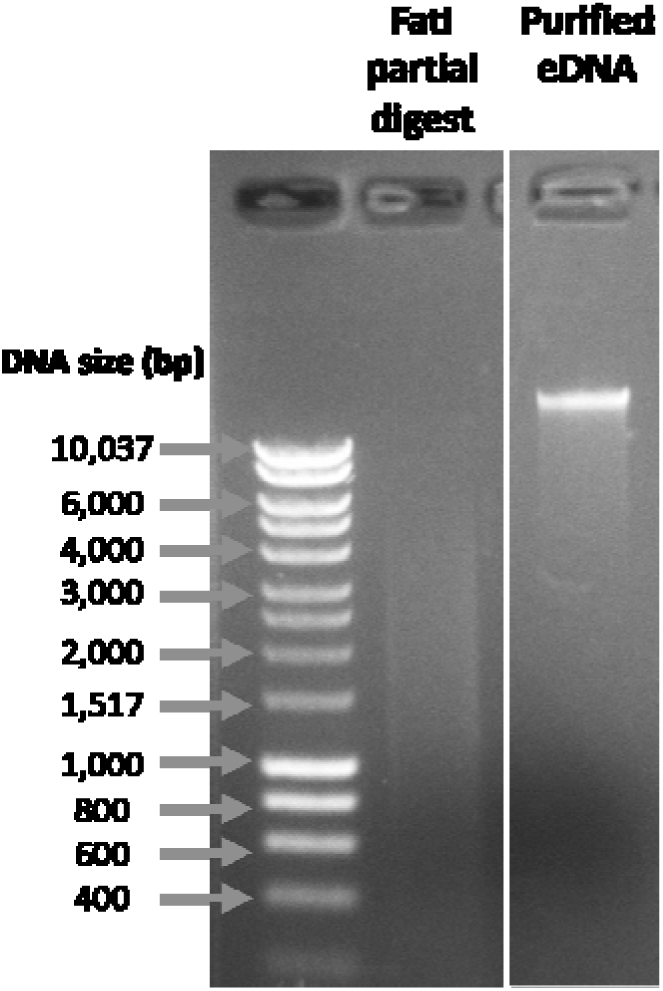
**Preparation of the FatI eDNA library.** eDNA extracted from soil was assessed by agarose gel electrophoresis (lane labelled “Purified eDNA”) then further purified by electroelution and partially digested with FatI restriction enzyme (stained aliquot in lane labelled “FatI partial digest”). Unstained DNA in the size range 0.6-1.4 kb from an aligned neighbouring lane was excised and gel-purified, then ligated into NcoI-treated pUCXMG to generate the final library.

**Supplementary Figure S5:**
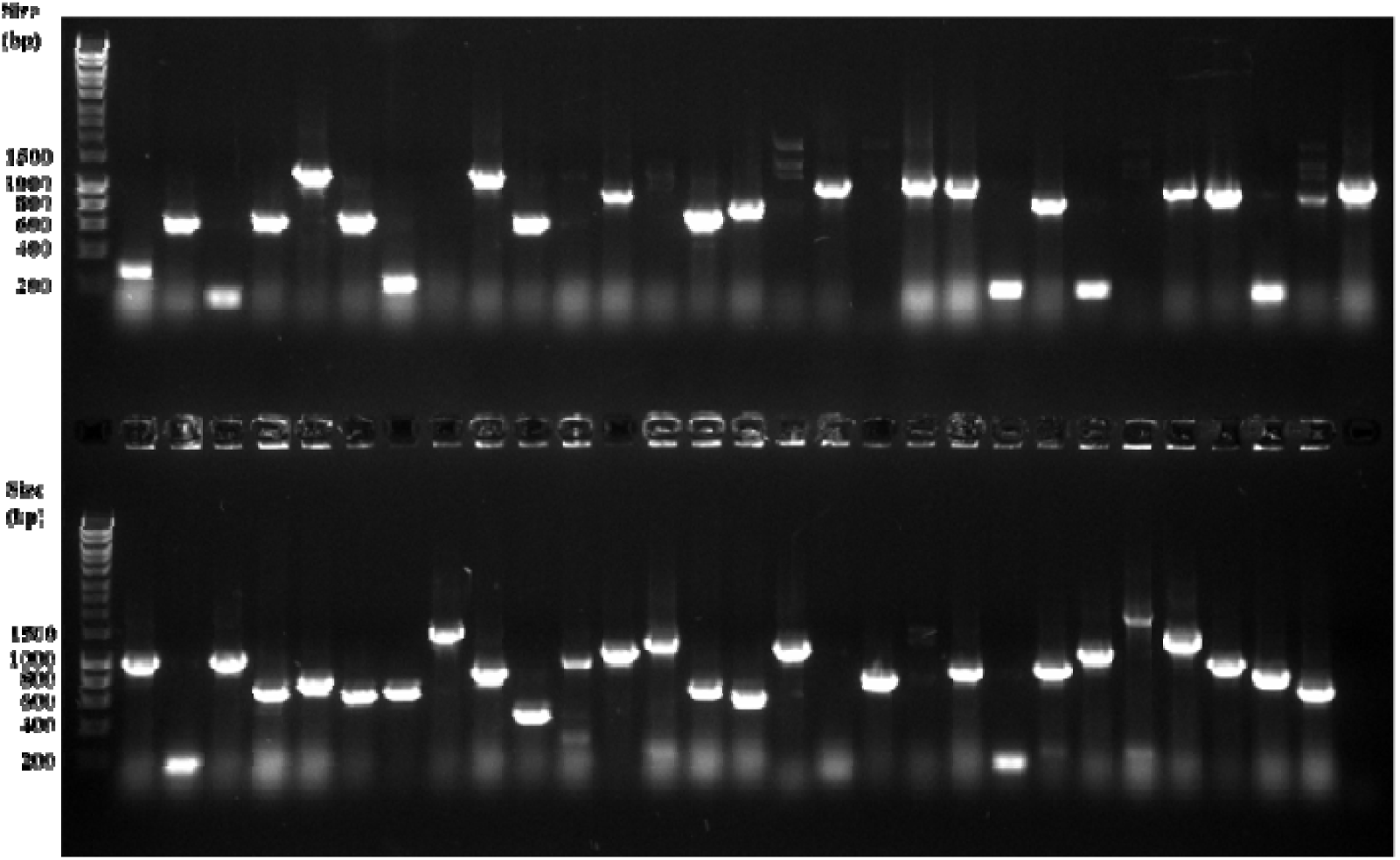
**Assessment of FatI eDNA library.** PCR colony screen of *E. coli* cells transformed with the pUCXMG-metagenomic DNA library ligation and plated on LB agar plates containing 20 µg/ml gentamycin. PCR was performed on 56 randomly-selected colonies using primers pUCX_for and pUCX_rev flanking the FatI/NcoI fusion site, yielding 49 amplicons in total (87.5%), with 41 of these being >500 bp (73.2%).

**Supplementary Table S1:** Annotation of ‘hit’ genes recovered from niclosamide selection of Swedish soil eDNA library

| Database description of best BLASTN match | Query coverage | E value |
| --- | --- | --- |
| efflux transporter outer membrane subunit [Betaproteobacter] | 85% | 1e-158 |
| multidrug efflux RND transporter [Prolixibacteraceae bacterium] | 74% | 3e-136 |
| multidrug efflux RND transporter [*Usitatibacter palustris*] | 78% | 1e-80 |

**Supplementary Table S2:** Zebrafish transgenic lines used in this study

| Transgene and allele | Reference |
| --- | --- |
| *Tg(14xUAS-E1B:NTR1.0-mCherry)c264* | Pisharath et al., 2007 |
| *Et(2xNRSE-Mmu.fos:KALTA4)gmc617* | Xie et al., 2012 |
| *Tg(5xUAS:tagYFP-P2A-MhqN1)jh542* | This work |
| *Tg(5xUAS:tagYFP-P2A-MhqN2)jh540* | This work |
| *Tg(5xUAS:tagYFP-P2A-MhqN3)jh545* | This work |
| *Tg(5xUAS:tagYFP-P2A-SagB1)jh543* | This work |

## Notes

### Competing Interest Statement

The authors have declared no competing interest.

