## Supplementary File S1 - Python Scripts for "A metagenomic library cloning strategy that promotes high-level expression of captured genes to enable efficient functional screening"

**Script 1**

**For each GenBank genome that we looked at, records that didn’t contain ‘plasmid’ in their description (i.e. only chromosome records) were selected for further analysis. For each chromosome, all features annotated as ‘CDS’ had their gene sequence and 3 bp immediately preceding the gene start extracted, and the frequency of various start codon and the nucleotide occurring at the -1 position (displayed as ()...; where () denotes the nucleotide in the -1 position and … represents the start codon nucleotides) were collated in a single row into an csv file. The genome length and GC content of each chromosome were also calculated and inputted into the same row along with the original genome file name and the record’s accession number.**

import os

from Bio import SeqIO

write_handle = open("Compilation_01.csv", "w")

write_handle.write("File,Accession,Description,Length,GC count,% GC,Total gene count,Total ATG start codon count,Total (C)ATG start codon count,Total (G)ATG start codon count,Total (A)ATG start codon count,Total (T)ATG start codon count,Total GTG start codon count,Total (C)GTG start codon count,Total (G)GTG start codon count,Total (A)GTG start codon count,Total (T)GTG start codon count,Total TTG start codon count,Total (C)TTG start codon count,Total (G)TTG start codon count,Total (A)TTG start codon count,Total (T)TTG start codon count\n")

for root, dirs, files in os.walk("."):

for filename in files:

if filename.endswith(".gbff"):

print (filename)

for gb_record in SeqIO.parse(open(filename, 'r'), "genbank"):

print (gb_record.description)

if "plasmid" not in gb_record.description:

total_gene_count=0

GC_count=0

percentage_gc=0

length=len(gb_record.seq)

description=gb_record.description

description="\"" + description + "\""

ATG_count=0

CATG_count=0

GATG_count=0

AATG_count=0

TATG_count=0

GTG_count=0

CGTG_count=0

GGTG_count=0

AGTG_count=0

TGTG_count=0

TTG_count=0

CTTG_count=0

GTTG_count=0

ATTG_count=0

TTTG_count=0

for n in range(0,len(gb_record.features)):

feature=gb_record.features[n] #Labels features

if feature.type=="CDS": #Only looking for CDS features

location=feature.location #Finds location of CDS within chromosome e.g. [100:300] (+)

start=location.start #Start location of CDS e.g. 100

end=location.end #End location of CDS e.g. 300

strand=location.strand #Strand - positive (forward) or negative (reverse)

if strand==1: #If CDS is located on forward strand of chromosome

gene=gb_record.seq[start-3:end] #extract entire CDS, including the 3 nts immediately preceding start codon

elif strand==-1: #If CDS is located on reverse strand of chromosome i.e. reverse complement

revcomp_gene=gb_record.seq[start:end+3] #extract entire CDS, including the 3 nts immediately following start codon (reversed)

gene=revcomp_gene.reverse_complement() #reverse complements gene so it reads in forward direction

total_gene_count+=1 #Adds up number of CDS sequences within file

if gene[3:6]=='ATG': #If start codon of gene is ATG

ATG_count+=1 #Adds up number of ATG start codons within CDS sequences

if gene[2]=='C': #If ATG start codon is immediately preceded by C i.e. FatI RE site

CATG_count+=1

elif gene[2]=='G':

GATG_count+=1

elif gene[2]=='A':

AATG_count+=1

elif gene[2]=='T':

TATG_count+=1

elif gene[3:6]=='GTG': #If start codon of gene is GTG

GTG_count+=1 #Adds up number of GTG start codons within CDS sequences

if gene[2]=='C': #If GTG start codon is immediately preceded by C

CGTG_count+=1

elif gene[2]=='G':

GGTG_count+=1

elif gene[2]=='A':

AGTG_count+=1

elif gene[2]=='T':

TGTG_count+=1

elif gene[3:6]=='TTG': #If start codon of gene is ATG

TTG_count+=1 #Adds up number of TTG start codons within CDS sequences

if gene[2]=='C': #If TTG start codon is immediately preceded by C

CTTG_count+=1

elif gene[2]=='G':

GTTG_count+=1

elif gene[2]=='A':

ATTG_count+=1

elif gene[2]=='T':

TTTG_count+=1

for n in gb_record.seq:

if n=='C' or n=='G':

GC_count+=1

percentage_gc=round(GC_count/length*100,2)

output = (filename + "," + gb_record.name + "," + description + "," +

str(length) + "," + str(GC_count) + "," + str(percentage_gc) + "," + str(total_gene_count) + "," +

str(ATG_count) + "," + str(CATG_count) + "," +

str(GATG_count) + "," + str(AATG_count) + "," + str(TATG_count) + "," +

str(GTG_count) + "," + str(CGTG_count) + "," +

str(GGTG_count) + "," + str(AGTG_count) + "," + str(TGTG_count) + "," +

str(TTG_count) + "," + str(CTTG_count) + "," +

str(GTTG_count) + "," + str(ATTG_count) + "," + str(TTTG_count) + "," +"\n")

write_handle.write(output)

print ("")

write_handle.close()

print ("Complete")

**Script 2**

**To combine data from multiple records (i.e. chromosomes) for the same bacterial genome, records with the same genome file name had their data merged into a single row and the accession numbers updated with a ‘+’ between the records combined.**

import csv

write_handle = open("Compilation_01_edited.csv", "w")

with open('Compilation_01.csv', newline='') as csvfile:

readfile = csv.reader(csvfile, delimiter=',')

old_list=list(readfile)

new_list=[]

for old_row in old_list:

new_list_files=[]

for new_row in new_list:

new_list_files+=[new_row[0]]

if old_row[0] not in new_list_files:

new_list.append(old_row)

elif old_row[0] in new_list_files:

#Join accession numbers

new_row[1]+=' + '+old_row[1]

#Join genome names

new_row[2]+=' + '+old_row[2]

#Add genome lengths

new_genome_length=int(new_row[3])

old_genome_length=int(old_row[3])

new_row[3]=str(new_genome_length+old_genome_length)

#Add C+G genome counts

new_gc_length=int(new_row[4])

old_gc_length=int(old_row[4])

new_row[4]=str(new_gc_length+old_gc_length)

#Recalculate genome GC percentage

genome_length=float(new_row[3])

gc_length=float(new_row[4])

new_gc_percent=round((gc_length/genome_length)*100,2)

new_row[5]=str(new_gc_percent)

#Add gene counts

new_gene_count=int(new_row[6])

old_gene_count=int(old_row[6])

new_row[6]=str(new_gene_count+old_gene_count)

#Add ATG starting genes

new_atg_count=int(new_row[7])

old_atg_count=int(old_row[7])

new_row[7]=str(new_atg_count+old_atg_count)

#Add (C)ATG starting genes

new_catg_count=int(new_row[8])

old_catg_count=int(old_row[8])

new_row[8]=str(new_catg_count+old_catg_count)

#Add (G)ATG starting genes

new_gatg_count=int(new_row[9])

old_gatg_count=int(old_row[9])

new_row[9]=str(new_gatg_count+old_gatg_count)

#Add (A)ATG starting genes

new_aatg_count=int(new_row[10])

old_aatg_count=int(old_row[10])

new_row[10]=str(new_aatg_count+old_aatg_count)

#Add (T)ATG starting genes

new_tatg_count=int(new_row[11])

old_tatg_count=int(old_row[11])

new_row[11]=str(new_tatg_count+old_tatg_count)

#Add GTG starting genes

new_gtg_count=int(new_row[12])

old_gtg_count=int(old_row[12])

new_row[12]=str(new_gtg_count+old_gtg_count)

#Add (C)GTG starting genes

new_cgtg_count=int(new_row[13])

old_cgtg_count=int(old_row[13])

new_row[13]=str(new_cgtg_count+old_cgtg_count)

#Add (G)GTG starting genes

new_ggtg_count=int(new_row[14])

old_ggtg_count=int(old_row[14])

new_row[14]=str(new_ggtg_count+old_ggtg_count)

#Add (A)GTG starting genes

new_agtg_count=int(new_row[15])

old_agtg_count=int(old_row[15])

new_row[15]=str(new_agtg_count+old_agtg_count)

#Add (T)GTG starting genes

new_tgtg_count=int(new_row[16])

old_tgtg_count=int(old_row[16])

new_row[16]=str(new_tgtg_count+old_tgtg_count)

#Add TTG starting genes

new_ttg_count=int(new_row[17])

old_ttg_count=int(old_row[17])

new_row[17]=str(new_ttg_count+old_ttg_count)

#Add (C)TTG starting genes

new_cttg_count=int(new_row[18])

old_cttg_count=int(old_row[18])

new_row[18]=str(new_cttg_count+old_cttg_count)

#Add (G)TTG starting genes

new_gttg_count=int(new_row[19])

old_gttg_count=int(old_row[19])

new_row[19]=str(new_gttg_count+old_gttg_count)

#Add (A)TTG starting genes

new_attg_count=int(new_row[20])

old_attg_count=int(old_row[20])

new_row[20]=str(new_attg_count+old_attg_count)

#Add (T)TTG starting genes

new_tttg_count=int(new_row[21])

old_tttg_count=int(old_row[21])

new_row[21]=str(new_tttg_count+old_tttg_count)

for new_row in new_list:

new_row[2]= "\"" + new_row[2] + "\""

new_row_string=','.join(new_row)

output=(new_row_string +"\n")

write_handle.write(output)

write_handle.close()
